# *Rhizoctonia solani* disease suppression: addition of keratin-rich soil amendment leads to functional shifts in soil microbial communities

**DOI:** 10.1101/2023.05.31.543058

**Authors:** Lina Russ, Beatriz Andreo Jimenez, Els Nijhuis, Joeke Postma

## Abstract

Promoting soil suppressiveness against soil borne pathogens could be a promising strategy to manage crop diseases. One way to increase pathogen suppression would be the addition of soil organic amendments, however the mechanism behind this effect remains unexplored. The presented study will focus on *Rhizoctonia solani* disease in sugar beet grown in two different soils. We aim to find how microbial communities and their molecular functions can be linked to *Rhizoctonia solani* disease suppression in sugar beet seedlings after soil is amended with a keratin-rich side stream from the farming industry. Amended soil samples were analyzed using shotgun metagenomics sequencing, and the disease score of plants infected with *Rhizoctonia* and grown in the same soil was collected. Results showed that both keratin-rich amended soils were rich in bacteria from the Flavobacteriaceae, Sphingobacteriaceae, Boseaceae, Phyllobacteriaceae, Caulobacteraceae, Oxalobacteraceae, Comamonadaceae, Rhodanobacteraceae and Steroidobacteraceae, as well as taxa from the phylum Bdellovibrionota, containing obligate predatory bacteria. The only fungal group that increased significantly was the Mortierellaceae family. Keratinases were abundant in the keratin-rich amended samples. Pfam domain enrichment analysis showed a decline in domains that could be annotated in both keratin-rich amended soils (Lisse ∼18% and Vredepeel ∼30%), showing an increase in unknown proteins. Among proteins that were enriched were those potentially involved in the production of secondary metabolites/antibiotics, proteins involved in motility, keratin-degradation, and contractile secretion system proteins (mostly type VI secretion system). These results could show that keratin-rich soil amendments can support the transformation into a disease suppressive soil by stimulating the same taxa that have been found in other disease suppressive soils. We hypothesize that these taxa are responsible for the suppression effect due to their genomic potential to produce antibiotics, secrete effectors via the contractile secretion system, and degrade oxalate, which is considered a virulence factor of *R. solani*, while simultaneously possessing the ability to metabolize keratin.

## Introduction

Food security is threatened by the increasing incidence of diseases in crops and the target to reduce the application of pesticides in 2030 by 50% following the new European Farm to Fork and Biodiversity strategies (Proposal on the sustainable use of plant protection products and amending Regulation (EU) 2021/2115). Enhancing soil suppressiveness against soil borne pathogens is a promising strategy to control diseases and crop losses caused by those pathogens. *Rhizoctonia solani* is the main causal agent of black scurf disease in potato and crown rot disease in sugar beet, among other arable crops. Disease suppression of *Rhizoctonia solani* in soils can be induced by two entirely different strategies. The first approach requires the successive planting of host crops in the presence of their pathogen resulting in reduced disease incidence over time. This phenomenon of disease decline by monocropping has first been described for take-all (*Gaeumannomyces graminis* var. tritici) in barley and wheat ((Gerlagh, 1968; Raaijmakers & Weller, 1998; Sarniguet et al., 1992). Take-all decline is defined as the spontaneous reduction in the incidence and severity of the disease and increase in yield occurring with continuous monoculture of the host crop following a severe attack of the disease (Schlatter et al., 2017). Also, for *R. solani*-induced diseases a decline has been reported in both field and pot experiments for several crops, i.e., wheat (Lucas et al., 1993; Mazzola & Gu, 2002; Roget, 1995; Wiseman et al., 1996), sugar beet (Expósito, 2017; Hyakumachi et al., 1990; Sayama et al., 2001), radish (Chern & Ko, 1989; Chet & Baker, 1980; Henis et al., 1978), potato (Jager & Velvis, 1995; Velvis et al., 1989) and cauliflower (Davik & Sundheim, 1984; Postma et al., 2010). The second approach to stimulate disease suppression requires the addition of organic amendments into soils. These amendments can be side streams from industrial food processing, farming, and agricultural activities. The re-use of such materials in agriculture aligns with the increasing interest in circularity, reducing the environmental impact and promoting valorization of waste products (Abbott et al., 2018; Alvarenga et al., 2017; De Corato, 2020). A large variety of organic products, residual streams from plant and animal production, food industry or society have been tested for their potential to suppress soilborne diseases. The effects are product- and disease dependent and therefore difficult to translate into practical applications so far (Bonanomi et al., 2010). The addition of compost ((Pérez-Piqueres et al., 2006; Termorshuizen et al., 2006; Tuitert et al., 1998) and cellulose-containing products (Clocchiatti et al., 2021; Kundu & Nandi, 1985) have been studied extensively, but effects were often unpredictable. Against *R. solani* particularly, the addition of chitin- and keratin-rich products have been shown to decrease disease (Andreo-Jimenez et al., 2021; Postma & Schilder, 2015). Disregarding the approach to induce *Rhizoctonia* suppressiveness in soils, several organisms or combinations thereof were commonly found to correlate with lower disease incidence in different crops and soils. These include members of the bacterial families *Oxalobacteriaceae, Comamonadaceae* and *Burkholderiaceae, Pseudomonadaceae* as well as members of the orders *Hyphomicrobiales* and *Sphingobacteriales* (notably *Flavobacterium, Chryseobacteria* and *Chitinophaga*) and the fungal family *Mortierellaceae* (Andreo-Jimenez et al., 2021; Bonanomi et al., 2010; Carrión et al., 2018; Carrión et al., 2019; Chapelle et al., 2016; Expósito, 2017; Gómez Expósito et al., 2017; Yin et al., 2021). A universal pattern of the responsible microorganisms and especially the underlying mechanisms for *Rhizoctonia* suppressive soils is still lacking, although several potential mechanisms of disease suppression have been proposed. One model attributes disease suppression to the expression of a non-ribosomal peptide synthetase of Pseudomonadaceae family members (Mendes et al., 2011). Another mechanism points to the role of oxalic and phenylacetic acid as the main driver of suppression. These organic acids induce a stress response in the specific rhizobacterial families which, during hyphal growth of *R. solani*, results in the onset of survival strategies including motility, biofilm formation and the production of secondary metabolites (Chapelle et al., 2016). These hypotheses have been proposed based on experimental set up in natural or agricultural soils without amendments. Therefore, disease suppression mechanisms via organic amendments remain unexplored.

The objective of the present work is to gain a deeper insight on how microbial communities and their molecular functions can be linked to *Rhizoctonia* disease suppression after the amendment with a keratin-rich side stream from the farming industry. We propose that the addition of keratinaceous compounds leads to an enrichment of specific genes in the microbial community that each by themselves have been shown to play a role in disease suppression.

## Material and Methods

### Selected samples and sequencing

Soil samples from two different fields located in Lisse and Vredepeel, The Netherlands, were collected one time in 2017. These soils differed in organic matter content (OM) and pH (resp. 1.5% OM, pH 7.2 and 4.0-4.3% OM, pH 5.5). Both soils were amended with a pig hair meal product (1.4 g/kg soil) (Darling Ingredients), hereafter referred to as keratin-rich amendment. Soil was amended with 1.2 g calcium nitrate (Ca(NO_3_)_2_ /kg soil was used as a control, to ensure the same nitrogen equivalents added as in the keratin-rich treatment (i.e., 0.2 g N/kg soil). All treatments were performed with 4 replicates and incubated for three weeks at room temperature. The total of 16 samples was then sampled for DNA extraction to perform shotgun metagenomics sequencing. The experimental set up, protocols and measurements are described in detail in (Andreo-Jimenez et al., 2021). Both soils were tested for damping off disease suppression caused by *Rhizoctonia solani* AG2-2IIIB in a sugar beet assay and were found to lead to a significant decrease in disease incidence (Andreo-Jimenez et al., 2021).

Libraries were prepared (Next Generation Sequencing Facilities, Wageningen University & Research, Wageningen, The Netherlands) and paired-end sequenced (2x150 nt) on Illumina NovaSeq 6000 platform (BaseClear B.V., Leiden, the Netherlands). The raw sequencing data used for this study are available on the NCBI sequence read archive (SRA) under BioProject number PRJNA966095 (reviewer link:

https://dataview.ncbi.nlm.nih.gov/object/PRJNA966095?reviewer=seaasdqnmti48epii65f2qfrm4.)

### Bioinformatics workflow

Raw reads were subjected to an all-in-one preprocessing using FASTP (Chen et al., 2018) with default settings for paired-end data. The remaining reads were passed to Kraken2 (Lu et al., 2022; Wood et al., 2019) for detection of taxa using a custom database. This database included complete bacteria (70813 genomes), complete archaea (739 genomes), complete fungi (1678 genomes), complete viral (14744 genomes) and complete protozoan (11151 genomes) databases. After building the database, a collection of soil microbes that have been identified as important players in this ecosystem previously (Andreo-Jimenez et al., 2021) and were either absent or underrepresented in the built database, were added to the library (Supplementary Table 1). Kraken2 was run in *--paired* mode on individual samples using the kraken2 standard output to be visualized with PAVIAN (Breitwieser & Salzberg, 2019). The absolute number of hits were extracted per family and genus level entry. These results were then used to investigate differential abundance distributions of microbial families and genera per soil with DESeq2 (Love et al., 2014) separately, using treatment (keratin vs control) as *design* input. Results were then visualized using ENHANCEDVOLCANO (Blighe et al., 2019).

**Table 1.** Sample description of Vredepeel and Lisse soils as well as sequencing, assembly, classification and clustering statistics per treatment and soil.

| sample ID | Year | Location | Treatment | # reads | # Plass-assembled proteins | Representative Protein clusters | Kraken2 classified (%) | Proteins with Pfam hits (%) |
| --- | --- | --- | --- | --- | --- | --- | --- | --- |
| 5_2_1 | 2017 | Lisse | control | 142978580 | 32591140 |  | 25.06 | 53.97 |
| 6_2_2 | 2017 | Lisse | control | 97003942 | 19979950 |  | 24.44 | 62.80 |
| 7_2_3 | 2017 | Lisse | control | 145297988 | 33944258 |  | 25.26 | 39.06 |
| 8_2_4 | 2017 | Lisse | control | 123152180 | 28018812 |  | 24.05 | 38.78 |
| Total |  |  |  |  | **114534160** | 64109105 |  |  |
| 45_12_1 | 2017 | Lisse | keratin-rich | 330315422 | 103987098 |  | 35.42 | 42.41 |
| 46_12_2 | 2017 | Lisse | keratin-rich | 87596756 | 19973315 |  | 37.01 | 43.13 |
| 47_12_3 | 2017 | Lisse | keratin-rich | 209939806 | 60993464 |  | 38.61 | 43.91 |
| 48_12_4 | 2017 | Lisse | keratin-rich | 160000448 | 41729783 |  | 36.64 | 30.07 |
| Total |  |  |  |  | **226683660** | 112892232 |  |  |
| 53_14_1 | 2017 | Vredepeel | control | 105381712 | 22595511 |  | 27.29 | 37.32 |
| 54_14_2 | 2017 | Vredepeel | control | 102527114 | 21888491 |  | 26.18 | 38.01 |
| 55_14_3 | 2017 | Vredepeel | control | 177871500 | 46720927 |  | 27.41 | 38.42 |
| 56_14_4 | 2017 | Vredepeel | control | 82682438 | 16963125 |  | 27.02 | 38.54 |
| Total |  |  |  |  | **108168054** | 78028651 |  |  |
| 93_24_1 | 2017 | Vredepeel | keratin-rich | 71538528 | 14684057 |  | 21.38 | 19.77 |
| 94_24_2 | 2017 | Vredepeel | keratin-rich | 104520536 | 23365019 |  | 21.32 | 19.86 |
| 95_24_3 | 2017 | Vredepeel | keratin-rich | 141049098 | 36036176 |  | 20.36 | 39.36 |
| 96_24_4 | 2017 | Vredepeel | keratin-rich | 120677294 | 29756912 |  | 20.96 | 19.81 |
| Total |  |  |  |  | **103842164** | 72610580 |  |  |

### Functional annotation and statistical analysis

Reads were assembled into proteins directly using the protein-level assembly tool PLASS (Steinegger et al., 2019) in paired-end mode to assemble QC-passed reads directly into proteins.

### Pfam enrichment analysis

For the enrichment analysis of protein domains in the dataset, the most recent Pfam annotations of Pfam-A and Pfam-N (15-11-2021 version) were downloaded from the EMBL-EBI FTP server (http://ftp.ebi.ac.uk/pub/databases/Pfam/) and converted into a *mmseqs* profile database (Steinegger & Söding, 2017). Using the protein assemblies of each sample separately as query, *mmseqs easy-search* was used to find Pfam profile hits for each sequence, choosing the best hits greedily (*--greedy-best-hits*). The hits per Pfam domain were counted and the resulting count matrix per sample was analyzed for differential abundance with DeSeq2 using “treatment” and “soil” as parameters in the design after prefiltering out low counts (< 50). PCA plots were created using log-ratio transformed output from the DESeq analysis. Pfam domains with a Log_2_Fold change of > 1 and P_adj_ < 0.01 were subset (Supplementary Table 2). Differentially abundant domains were plotted using EnhancedVolcano (FCCutoff= 1.0, pCutoff= 10e-6).

### Detection and classification of keratinases

The Peptidase Full-length Sequences were downloaded from the MEROPS peptidase database version 12.1 (Rawlings et al., 2017) and converted into a DIAMOND database (Buchfink et al., 2021; Buchfink et al., 2015). Assembled proteins of keratin-amended soils from Lisse and Vredepeel were searched against the peptidase database with a DIAMOND *blastp* run with a block size value of 12 and only showing the single best hit and further default settings. Protein sequences of the positives hits were extracted, and taxonomic information was added by running *mmseqs taxonomy* using the UniRef100 database as a reference with default settings. Information on the MEROPS family, the taxonomic lineage, the soil, and the sum of hits of each family per MEROPS family was combined using custom scripts in python and R and visualized using ggplot2. For the sake of visibility, a subset was taken containing known keratinases (J. Qiu et al., 2020) and microbial families that were shown to be more abundant in amended soils. These data were also visualized using ggplot2.

### Untargeted approach using protein clustering

Due to the extremely high number of reads that could not be assigned to any taxonomic group, we also performed an untargeted approach to be able to classify proteins unique to the treatment independent from homologies with entries in available databases. To this end, protein assemblies were combined into treated and control samples per soil. The data load was downsized by clustering proteins and only keeping the representative sequences: We first converted the protein assemblies into *mmseqs* databases and used *linclust* of the mmseqs2 package on the pooled assemblies of the control and the treated samples in bidirectional coverage mode with sequence identity threshold and an alignment coverage of 80% (--min-seq-id 0.8 --cov-mode 0 -c 0.8) to preserve the multi domain structure of proteins. Representative proteins per cluster were then extracted from the clustering results (*mmseqs createsubdb* inDB_clu inDB inDB_clu_rep) and converted into a fasta file (*mmseqs convert2fasta* inDB_clu_rep inDB_clu_rep.fasta). The representative proteins of the control per soil were converted into a DIAMOND database and used as query against the representative proteins of the keratin-treated samples in a *diamond blastp* run with a block size value of 12 and only showing the single best hit and further default settings. Unaligned proteins were stored and used to create an mmseqs2 database to identify taxonomy and functions of proteins present using the *mmseqs taxonomy* against the Uniref100 database. The result of the functional characterization of proteins was exported into a .tsv file also containing the taxonomic lineage. Next to that we performed a functional annotation using the standalone version of KofamScan (version 1.3.0) with the standard database (KO profiles (release 24-Apr-2023) KO list (release 26-Apr-2023)) (Aramaki et al., 2019). The datasets were then merged into a single file per soil containing functions and taxonomy in different columns. The table was parsed and families with a total of more than 200 protein hits were extracted. From those we used the top 100 most abundant KO identifiers and plotted the data using ggplot2.

## Results

### Sequencing results, annotation, and community profile

A summary of the samples and sequenced reads per sample is available in Table 1. Samples from Lisse soil were sequenced much deeper than Vredepeel soil samples. The amount of reads that could be classified using the customized Kraken database was low and depended on the soil and the treatment it received (Table 1). Interestingly, the percentage of classified reads in Lisse soils that had been supplemented with the keratin-rich amendment increased from 24.70±0.46 to 36.92±0.89, whereas classified reads in Vredepeel soil dropped after having received keratin-rich treatment (from 26.98±0.40 to 21.01±0.35). Most of all classified reads disregarding soil and treatment could be attributed to bacteria (96.81% - 98.59%), followed by fungi (0.16-0.31%). Viral and protozoan contribution to the metagenome was less than 0.01% of the total reads.

### Taxonomic shifts after keratin-rich amendment in soil

As bacterial and fungal taxa represented the largest groups in the identifiable fraction, we focused our further analysis on these. Differential abundance analysis of classified taxa on family level showed a similar shift in both soils upon the addition of the keratin-rich product. Both soils increased in *Flavobacteriaceae* and *Sphingobacteriaceae* (*Bacteriodetes*), *Boseaceae, Phyllobacteriaceae* and *Caulobacteraceae* (*Alphaproteobacteria*), *Oxalobacteraceae* and *Comamonadaceae* (*Betaproteobacteria*), *Rhodanobacteraceae* and *Steroidobacteraceae* (*Gammaproteobacteria*), as well as one (Vredepeel; *Bacteriovoraceae*) or even three (Lisse; *Bacteriovoraceae, Bdellovibrionceae, Halobacteriovoraceae*) families of the bacterial phylum *Bdellovibrionota*, containing obligate predatory bacteria (Fig. 1A & B). The only fungal group that increased significantly was the zygote fungal family of *Mortierellaceae* (Fig. 1B. Differences between the two soils were represented by *Nitrosomanadaceae, Rhizobiaceae* and *Bdellovibrionaceae* which did show a significant increase after keratin-rich amendment in Lisse soil, but not in Vredepeel soil. Furthermore, *Micrococcaceae* and *Caulobacteraceae* were more abundant in Lisse soil and did show a steeper increase after keratin amendment (Fig. 1A).

**Figure 1.**
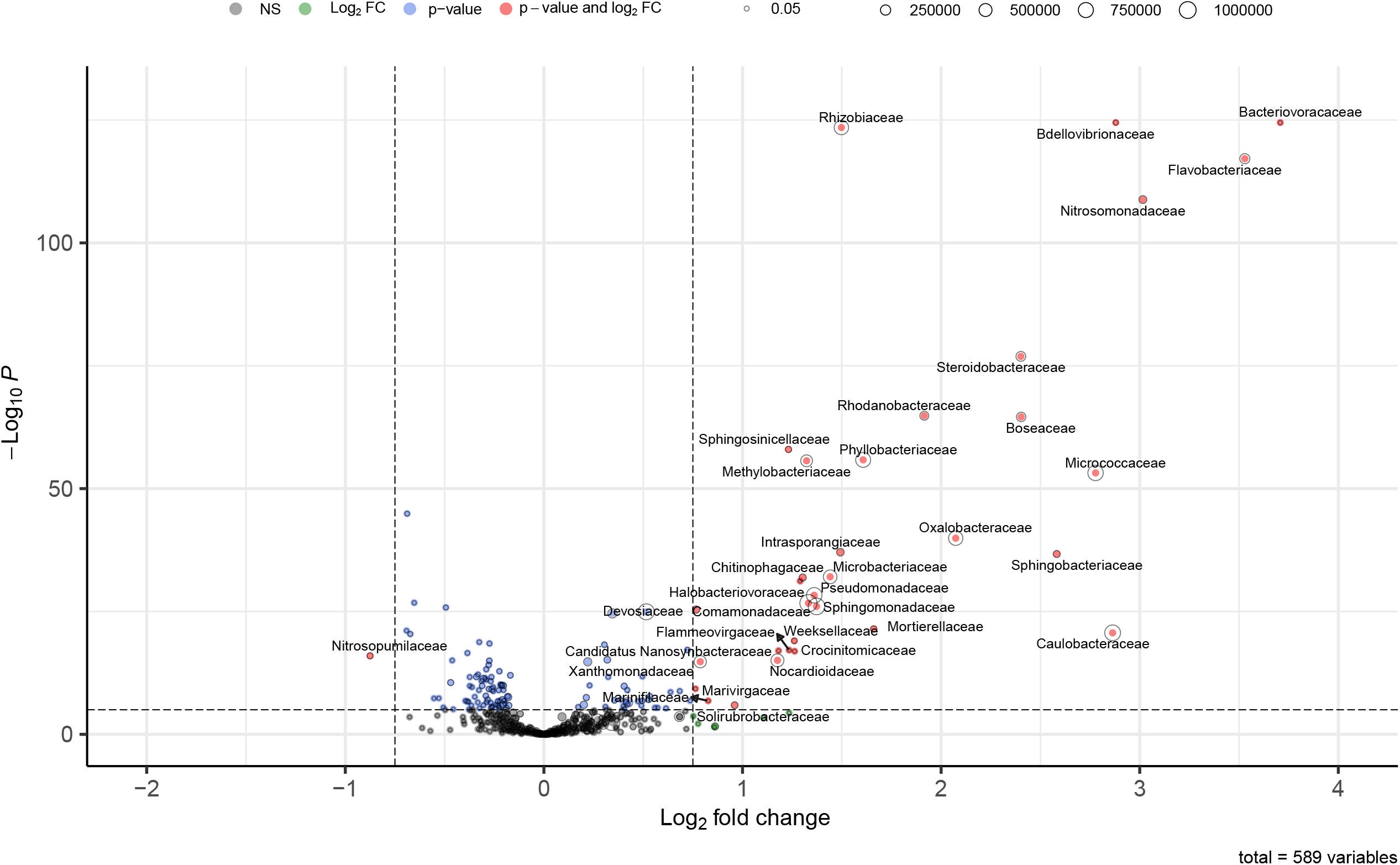
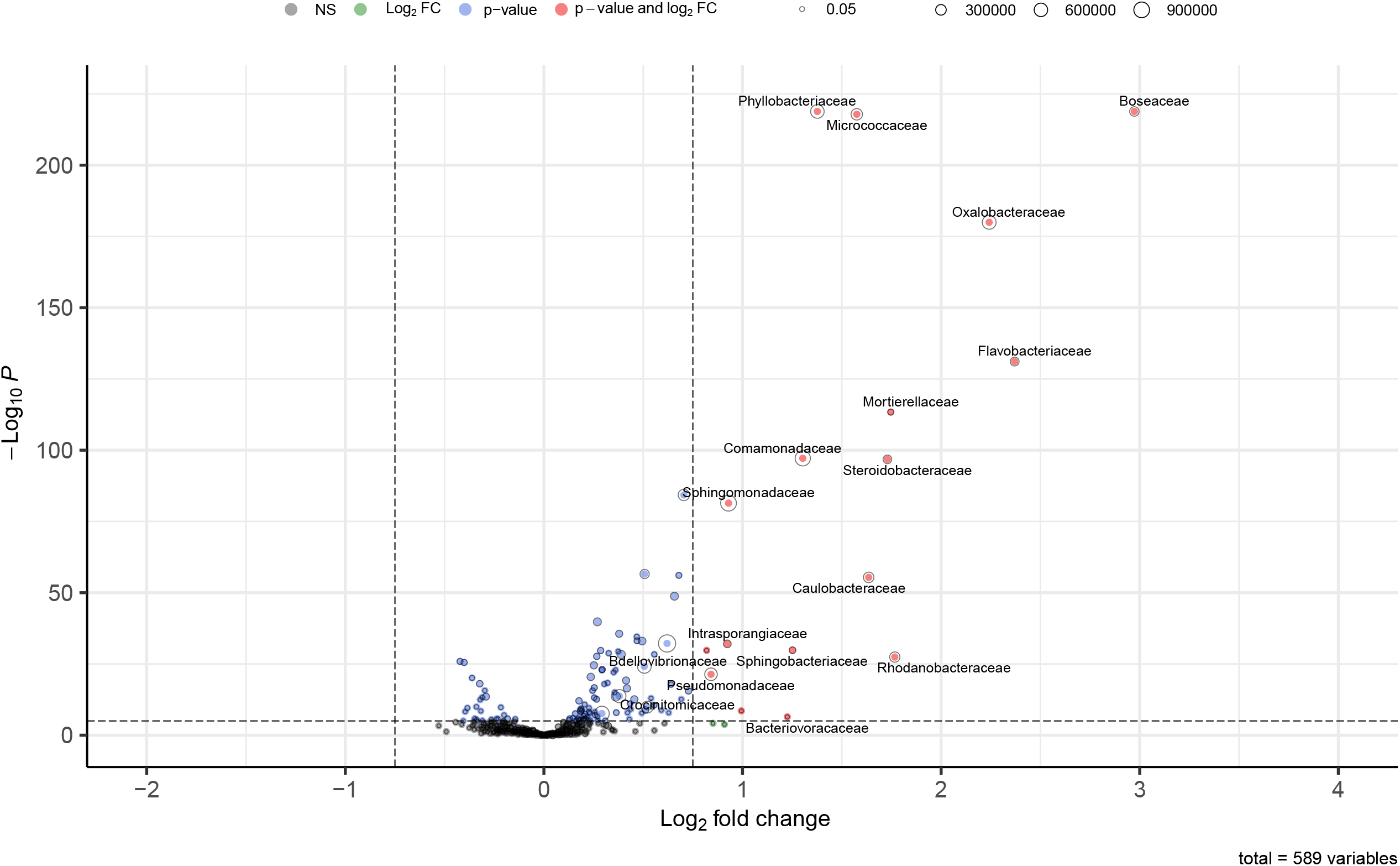
Differentially abundant microbial families after keratin-rich amendment. Volcano plots represent bacteria and fungi, which are differentially abundant within keratin-rich product treatment compared with the calcium nitrate control, in Lisse (a) and Vredepeel (b) soils (FDR=1x10^−6^, Log_2_FC=0.75). Based on normalized reads obtained after the Kraken2 workflow. Bubble size depicts the abundance of the family as the total number of reads.

### Keratinolytic potential of abundant taxa

Amending both soils with a keratin-rich compound compared to an inorganic source of nitrogen (Ca(NO_3_)_2_) led to similar changes in the taxonomic composition of the identifiable microbial fraction after an incubation time of three weeks. It was therefore expected that the potential to degrade keratinaceous compounds could play a role in the enrichment of certain taxa. The taxonomic families shown to be more abundant after the incubation with keratin showed a high occurrence of proteases belonging to MEROPS families of known keratinases (Fig. 2). Lisse soil did contain a much higher number of protease matches. However, differences between Lisse and Vredepeel soil response are difficult to interpret due to the uneven read depth of both samples. The absence of a certain protein family could also be caused by insufficient depth, especially of organisms with a lower abundance. In Lisse soil proteases of all known keratinase families (REF) could be recovered for *Steroidobacteraceae, Solirubrobacteraceae, Rhodanobacteraceae, Pseudomonadaceae, Oxalobacteriaceae, Nocardioidaceae, Nitrosomonadaceae, Comamonadaceae* and *Chitinophagaceae*. The MEROPS families with the highest number of hits were S09 (24.4% of subset hits), M04 (15.9%), M38 (14.1%) and S01 (10.4%). Much less common in frequency as well as in taxonomic distribution were the metalloprotease families M36 (0.14%) and M32 (0.13%).

**Figure 2.**
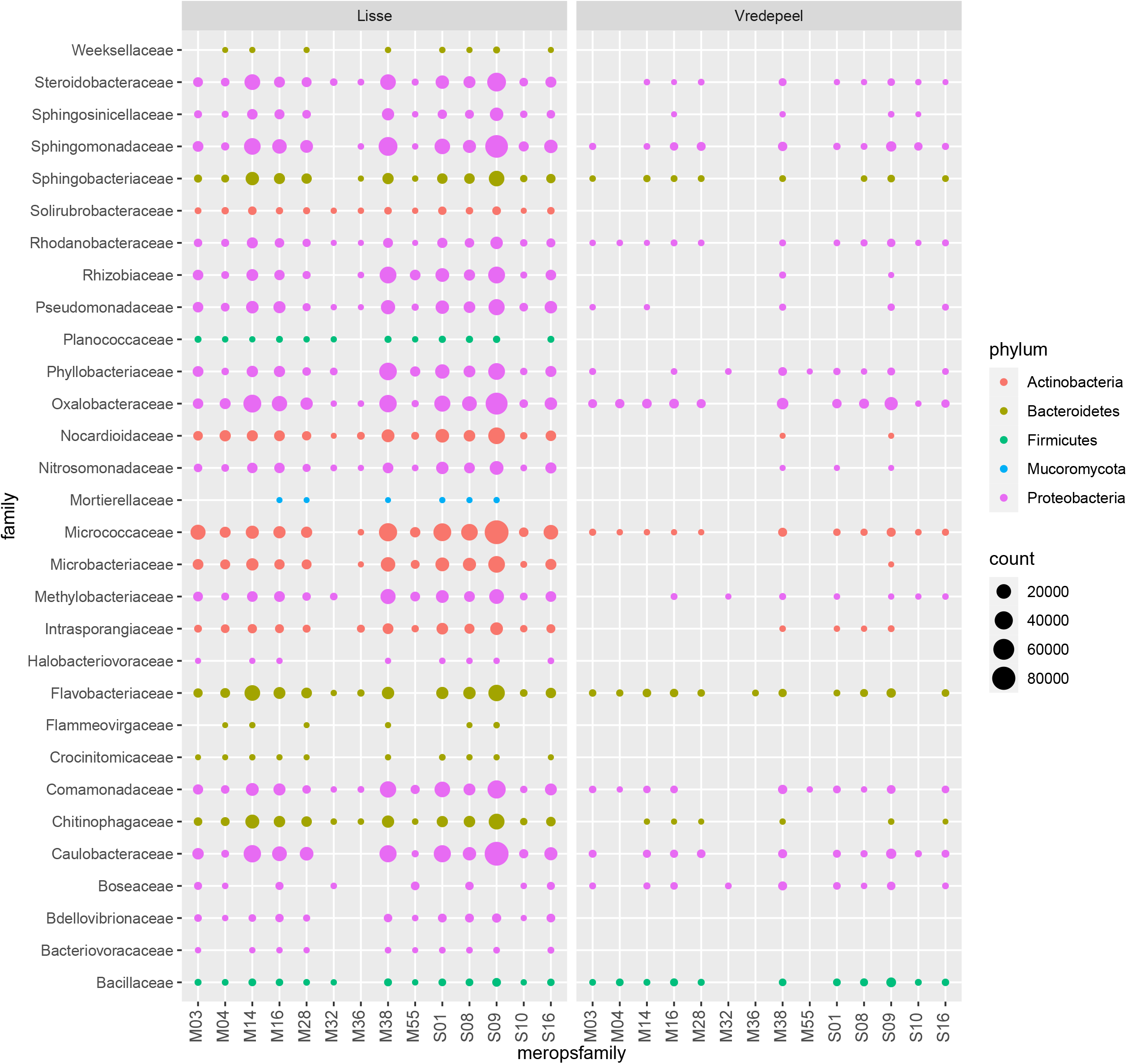
Counts of hits on MEROPS protein families with known keratinolytic representatives within microbial families more abundant in soils having received a keratin-rich amendment.

### Specific protein domains were enriched after keratin-rich amendment

Next to the taxonomic classification of reads and keratinase-like protease families, a classification on Pfam domains of assembled proteins was used as a proxy for functional changes in the microbiome upon the soil amendment with a keratin-rich product. The fraction of proteins carrying a Pfam domain found in the database differed between the control and the keratin-amended soils. If soils had received a keratin-rich amendment the number of proteins that could be assigned a Pfam domain dropped in both soils (Tab. 1) (Lisse 39.88±5.69% with keratin-rich amendment, 48.65±10.22% in control; Vredepeel 24.70±8.46% with keratin-rich amendment, 38.07±0.48% in control) although standard deviation was high. The biological replicates of each treatment category per soil did show similar Pfam patterns and formed distinct clusters in a Principal component analysis (PCA) (Fig. 3). The keratin-rich amendment did have an influence on the protein domain abundances, resulting in a shift along PC1 that was visible in both soils (Fig. 3).

**Figure 3.**
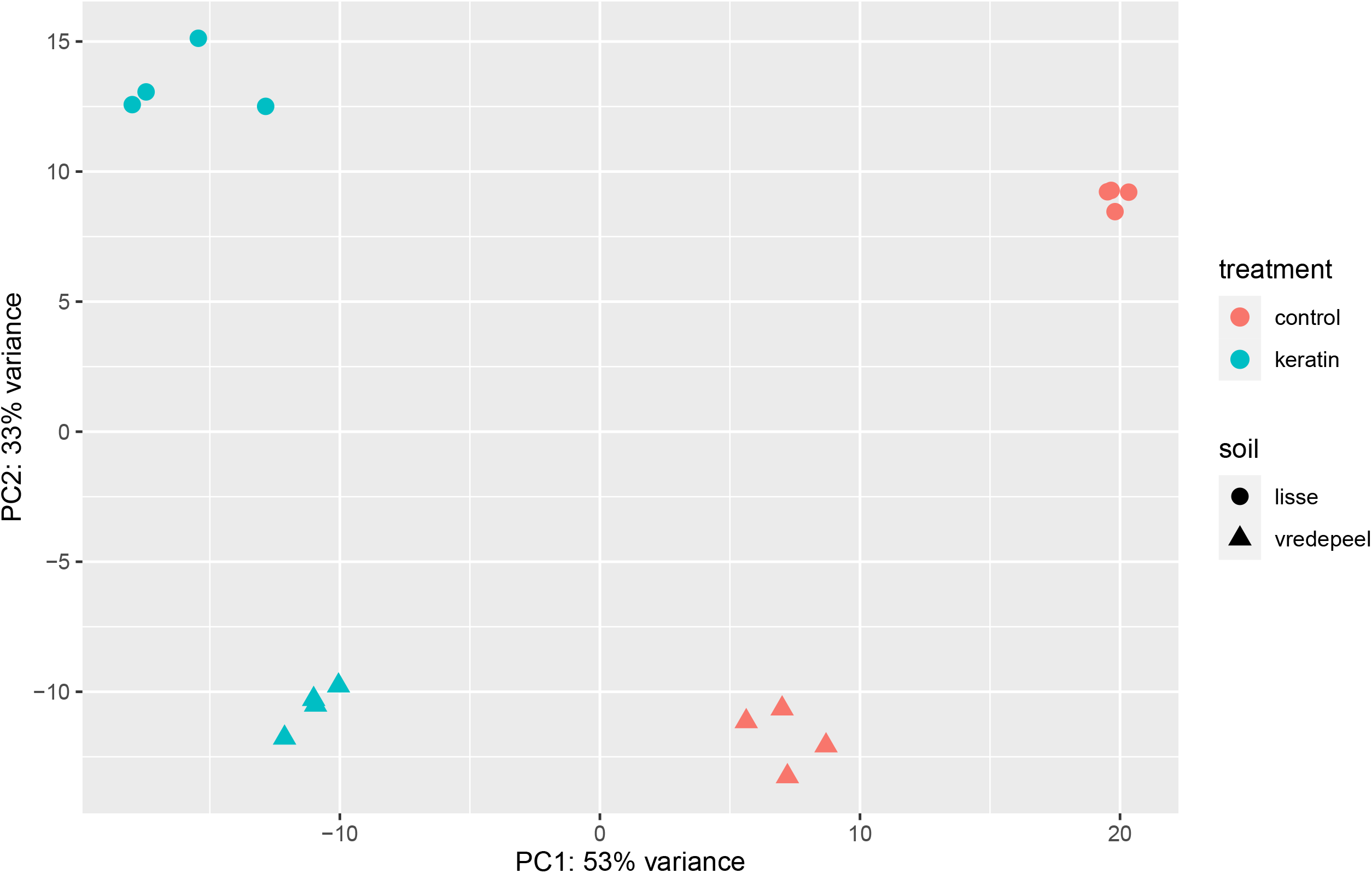
Principal component analysis (PCA) of Pfam domain composition of keratin-rich and control treatment in Lisse and Vredepeel soil.

In Vredepeel soil 9350 Pfam domains were detected that had a nonzero total read count, 2209 of them having a P adjusted value of < 0.01. In Lisse soil, 10133 different protein domains could be detected, of which 4248 had a P adjusted < 0.01. To identify which domains were enriched if the keratin-rich product was added to the soil, a cutoff of Log_2_FC > 1.0 was chosen in addition to a P adjusted < 0.01 (Supplementary Table 2). Results showed that 480 protein domains were enriched in both Lisse and Vredepeel soils after having received an amendment with keratin. An additional 1227 and 179 Pfam domains were exclusively enriched in Lisse and Vredepeel soil respectively. Of the 480 Pfam domains enriched in both types of soil after keratin-rich amendment 168 were associated with domains of unknown functions (DUFs) and could not be characterized further. Potential domains of interest included a domain related to chitinase C (PF06483), several components of the type VI secretion system (T6SS) (PF18443, PF18426, PF13296, PF06744), domains related to the proteins involved in the biogenesis of bacterial cell appendages (PF00419, PF02753, PF06864, PF09977, PF10671, PF13681, PF15976, PF16823, PF16970) and flagella (PF08345, PF02108, PF03646, PF03961, PF05247, PF05400, PF06490, PF07196, PF07317, PF10768), peptidases with a domain similar to the M9 family (PF01752, PF08453), enhancin-like metallopeptidase domains of family M60 (PF17291, PF13402), microbial collagenases (PF01752, PF08453) and a number of phage(-like) proteins (PF00959, PF03374, PF04233, PF04466, PF05065, PF05133, PF05367, PF05954, PF06761, PF06892, PF09306, PF09669, PF13876, PF14395, PF16083, PF16510, PF18013, PF18352).

### Functional shifts in disease suppressive soils

To identify proteins that were unique to keratin-enriched amendment regardless of the soil, we clustered all proteins per soil and treatment resulting in 64109105 and 78028651 for Lisse and Vredepeel soils having received the control treatment and 112892232 and 72610580 for the ones amended with a keratin-rich substrate respectively (Tab.1). After alignment of clustered protein representatives of the control to the keratin-rich treatment using diamond and keeping only a single best hit, 554610 proteins remained unaligned. These were unique to keratin-rich amendment. As downstream processing required the functional annotation of those proteins, KofamScan was used to assign KEGG orthologies. Of all unique proteins only 53435 (3.44%) could be assigned. Overall, there was a strong overlap in functional groups with Pfam enrichment analysis despite the different approach (Fig.4). This included proteins potentially involved in the production of secondary metabolites/antibiotics such as 4’-demethylrebeccamycin synthase (K19888) found in representatives of the *Burkholderiales, Sphingomonadaceae* and several *Actinomycetes*, as well as gramicidin S synthase 2 (K16097) predominantly found in *Bacteroidota*. Proteins that could play a role in keratin degradation, such as serine protease (K14645) present in many organisms and the collagenase kumamolisin (K08677) found in *Oxalobacteraceae, Rhodanobacteraceae, Bacillaceae, Micrococcaceae* and *Intersporangiaceae* as well as several transporters potentially involved in transport of keratin and/or its derivatives such as a basic amino acid/polyamine antiporter (K03294). Interesting were also several enzyme families that degrade more complex substrates such as hyaluronate lyase (K011727), alpha-L-rhamnosidase (K05989) and arabidan endo-1,5-alpha-L-arabinosidase (K06113), 2,6-dioxo-6-phenylhexa-3-enoate/TCOA hydrolase (K22677) or exo-acting protein-alpha-N-acetylgalactosaminidase (K25767) and corresponding transporters like cholesterol transport system auxiliary component (K18480). In addition, several major facilitator superfamily transporters and multidrug resistance proteins were recovered from soils amended with keratin-rich amendments (K03762, K05519, K08166, K18353, K18555, K18904, K18926), a trait shared among a wider range of microorganisms. The presence of a hemoglobin/transferrin/lactoferrin receptor protein (K16087) seems to be a trait of *Pseudomonadota*. Several genes involved in the acquisition of iron could be identified such as a ferric enterobactin receptor (K19611) with the highest number of hits found in *Bacteriodota, Caulobacteraceae, Sphingomondaceae* and *Oxalobacteraceae* and an iron−siderophore transport system substrate−binding protein (K25109), which seems to be more widely distributed. An interesting candidate protein is an uncharacterized protein (K07126) especially enriched among *Oxalobacteraceae, Phyllobacteriaceae, Comanomadaceae, Pseudomonadaceae* and the only protein found in the fungal family of *Mortierellaceae* at the set cut-off. Other interesting features include proteins involved in motility (K02416, K02397, K10564) and the type VI secretion system secreted protein VgrG (K11904) and type VI secretion system protein ImpH (K11895).

**Figure 4.**
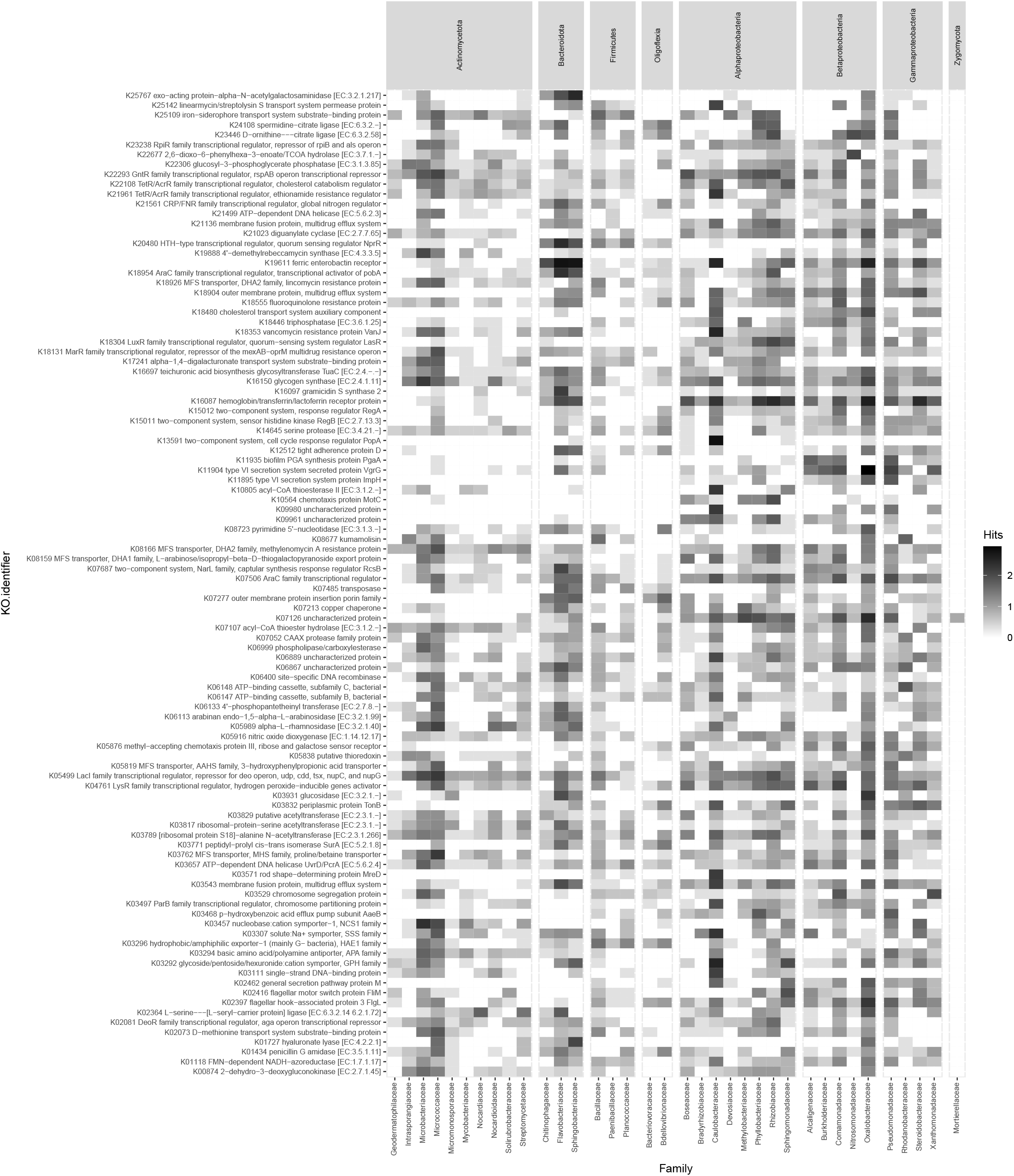
Abundance of the top 100 most abundant KEGG orthologies in microbial families with a total of ≥200 protein hits. The number of protein hits was Log-transformed Log10(n+1) and families grouped by a higher taxonomic order (Phylum; *Pseudomonadota* split on the level of class).

## Discussion

In the presented study we showed that keratin-rich products amended to the soil can affect the abundance of specific bacterial and fungal families in both soils studied. Microbial families that changed most significantly three weeks after the amendment of a keratin-rich substrate were *Flavobacteriaceae* and *Sphingobacteriaceae* (Bacteriodetes), *Boseaceae, Phyllobacteriaceae* and *Caulobacteraceae* (Alphaproteobacteria), *Oxalobacteraceae* and *Comamonadaceae* (Betaproteobacteria), *Rhodanobacteraceae* and *Steroidobacteraceae* (Gammaproteobacteria), as well the family of obligate bacterial predators *Bacteriovoraceae*. Within the fungal kingdom only the *Mortierellaceae* increased upon the addition of keratin. Previous work using metabarcoding analysis on the same soil samples revealed also *Oxalobacteraceae, Flavobacteraceae* and *Mortierellaceae* as the microbial families that were associated with the most pathogen-suppressive treatments i.e., keratin and chitin-rich amendments (Andreo-Jimenez et al., 2021). We assume that the first enrichment of taxa and functions is due to the addition of the organic amendment: Microorganisms that are able to degrade keratin-rich materials directly or those able to metabolize degradation products can propagate. The high abundance of potential keratinolytic enzymes in taxa that increased upon the addition of the keratin-rich amendment supports this idea. Despite the different nature of keratin (polypeptide) and chitin (polysaccharide) molecules, their amendment to soil seems to result in a comparable shift in the soil microbiome as was shown by Andreo-Jimenez et al. (2021). Wieczorek et al. (Wieczorek et al., 2019) confirmed that members of the *Oxalobacteraceae* and several *Bacteroidetes* families were the initial chitin degraders in agricultural soils using stable isotope probing. Interestingly, they could also show an increased labelling in bacterial predators, suggesting that microorganisms that degrade chitin are prey of *Bacteriovoraceae* and *Bdellovibrionaceae*. We assume that the increase of these two families in our experiment has the same reason. Keratin-rich and chitin-rich amendment stimulate similar degraders and therefore also similar predators.

As the complete degradation of keratin, similarly to chitin and cellulose, requires the synergistic action of several enzymes it is impossible to deduce how the breakdown of keratin is orchestrated in the group microorganisms (Jingwen Qiu et al., 2020). What does the succession look like or are some community members able to perform the complete degradation pathway alone or only as a group are questions not to be answered with the data available. Comparing more enriched taxa in amendment-induced *R. solani* suppressive soils from this study and Andreo-Jimenez et al. (Andreo-Jimenez et al., 2021) with those from successive monocropping strategies resulted in comparable microbiome patterns including a higher abundance of *Pseudomonadaceae, Sphingomonadaceae, Burkholderiaceae, Xanthomonadales, Oxalobacteraceae, Caulobacteraceae, Sphingobacteriaceae, Chitinophagaceae* and *Flavobacteriaceae* in disease suppressive soils (Chapelle et al., 2016; Cordovez et al., 2015; Expósito, 2017; Mendes et al., 2011; Yin et al., 2021). It seems that selected organic amendments can stimulate the same groups that are associated with disease suppression without the addition of an amendment. This implies that the ability of a microorganism to degrade keratin, chitin and/or their derivates indirectly contributes to the transformation of a conducive into a *Rhizoctonia-*suppressive soil. Knowledge on the functional mechanism behind this phenomenon however remains patchy, mostly restricted to certain single taxa. Carrion et al. (2018) could show that the production of sulfurous volatile compounds with antifungal activity by *Burkholderiaceae* lead to disease suppression of *R. solani* in situ (Carrión et al., 2018). They could also show that members of the *Chitinophagaceae* and *Flavobacteriaceae* in the root endosphere were enriched in disease-tolerant plants (Carrión et al., 2019). These families encoded enzymes with enhanced enzymatic activities associated with fungal cell-wall degradation and secondary metabolite biosynthesis being a direct or indirect protective trait. Mendes *et al*. attributed disease suppression to the expression of a nonribosomal peptide synthetase of *Pseudomonadaceae* (Mendes et al., 2011). Chapelle *et al*. propose a vital role for oxalic and phenylacetic acid which, during hyphal growth of *R. solani*, induce a stress response of specific rhizobacterial families leading to the onset of survival strategies including motility, biofilm formation and the production of secondary metabolites (Chapelle et al., 2016).

We hypothesize that certain groups of microorganisms that are increasing in abundance after the addition of the keratin-rich amendment, due to their ability to metabolize the substrate (as was shown on their repertoire of relevant proteases), also encode versatile functions that are commonly associated with disease suppression. Among these are the contractile injection systems, such as the type VI secretion system (T6SS) or the extracellular contractile injection system (eCIS). Both are nanomachines resembling the bacteriophage puncturing structure, to secrete a variety of effectors that play a significant role in competition. These effectors can be injected into neighboring bacterial or eukaryotic cells or the environment causing arrest of their growth or even cell death, scavenging of nutrients or being used for cell to cell signaling (Coulthurst, 2019; Gallegos-Monterrosa & Coulthurst, 2021; Geller et al., 2021). Several effectors secreted via this system have even been described to have a direct antifungal effect (Trunk et al., 2019; Trunk et al., 2018). It is a common feature in *Pseudomonadota*, where more than 25% of sequenced genomes are estimated to carry at least one T6SS (Boyer et al., 2009) and it is especially abundant in plant-associated microbes (Bernal et al., 2018). Our findings showed that proteins belonging to contractile injection systems are enriched in the metagenome after keratin-rich amendment, especially in β- and γ-Proteobacteria.

The ability to produce antibiotics or bacteriocins is a feature often associated with disease suppression. Although it was not possible to identify complete biosynthetic gene clusters from the data available, due to the lack of genetic context because of protein-level assembly, we could identify a few proteins that might play a role in antibiosis, such as 4’-demethylrebeccamycin synthase or gramicidin S synthetase 2. From literature it is known that many representatives of enriched taxa have been described for their ability to produce antimicrobial compounds: Several members of the *Oxalobacteraceae*, among which *Massilia, Duganella, Pseudoduganella* and *Janthinobacterium* are able to produce several different antimicrobial compounds. The best studied being the purple pigment violacein, a bisindole with antimicrobial activity among other biological functions (Seong Yeol Choi et al., 2015; S. Y. Choi et al., 2015; Dahal et al., 2021). The phylum of *Actinomycetota* is known as a group harboring a wealth of gene clusters encoding antimicrobial agents. Most antibiotics in clinical use are originally isolated from these microbes (Genilloud, 2017; van der Meij et al., 2017). And within the *Bacteroidota* a recently published study has identified a high density of biosynthetic gene clusters per genome, with the genus *Chitinophaga* as the most interesting in terms of abundance and diversity (Brinkmann et al., 2022). This aligns with the discovery of Carrion *et al*. who found a consortium of endophytic *Chitinophaga* and *Flavobacterium* to consistently suppress fungal root disease, by the expression of chitinases, nonribosomal peptide synthases and polyketide synthases (Carrión et al., 2019).

Oxalotrophy, the ability to use oxalic acid as a carbon source, is a rare trait in bacteria, restricted to a few representatives of the phyla *Actinobacteria, Firmicutes* and *Pseudomonadota*. It is however often found in microorganisms that are associated with plants (Hervé et al., 2016). A study on *Burkholderia* strains in the rhizosphere of white lupin showed that 98% of the strains were able to grow on plant-secreted oxalate as a carbon source compared to only 2% of other strains isolated from the same environment. Oxalic acid is therefore suggested to stimulate the recruitment of plant-beneficial members from the soil microbiome (Kost et al., 2014). *Rhizoctonia solani*, like many other pathogenic fungi, is also able to produce oxalic acid to acidify host tissue and sequester calcium from host cell walls (Dutton & Evans, 1996; Palmieri et al., 2019; Yang et al., 1993). Increased virulence of *R. solani* has been shown to coincide with elevated levels of oxalic acid being excreted (Nagarajkumar et al., 2005). This leads Chapelle et al. to propose that the stress response activated under attack by the oxalic and phenylacetic acid produced either by *R. solani* itself or released from plant roots shifts leads to a *Rhizoctonia*-suppressive microbiome (Chapelle et al., 2016). Taxa that are also found to be enriched after keratin-rich amendment, especially members of the *Oxalobacteraceae* are known to metabolize oxalate and could thereby influence the invasion success of *R. solani* or detoxify oxalate for other community members.

We realize that many questions remain, such as the role of the major fraction of unknown proteins and/or organisms that were increased upon keratin-rich amendment. That it is currently not possible to consider that ‘microbial dark matter’ is a problem, because it could have significant implications on the mechanism of disease suppression of soils. In addition, the role of *Mortierella* and its relatives remains elusive for now. It has been shown for several fungal species that the ability to degrade keratin often co-occurs with the ability to break down other recalcitrant substrates such as cellulose and chitin via lytic polysaccharide monoxygenases (Lange et al., 2016). That this fungal group is enriched after keratin-rich amendment is due to its ability to degrade complex compounds therefore suggests itself. Interestingly, it has been shown recently that *Mortierella* species harbor bacterial endophytes of the *Burkholderiaceae* family (Takashima et al., 2018), some of which even protect the fungal host from nematode attack by the production of a biosurfactant (symbiosin) with antibiotic properties (Büttner et al., 2021; Büttner et al., 2023).

Taken together we propose that the amendment of a keratin-rich side stream product from the farming industry (but probably also chitin and cellulose) stimulates the enrichment of taxa that have been associated with *Rhizoctonia* disease suppression previously. Many of these taxa can metabolize the substrate or its derivatives and are well equipped for a life in the rhizosphere that could explain their contribution to disease suppression. This includes, among others, the ability to produce a wide array of secondary metabolites and effectors that can be secreted via contractile injection systems and the ability to use oxalic acid.

## Acknowledgements

We would like to thank Stefan Aanstoot, Carin Lombaers and Mirjam Schilder for technical assistance. This research was funded by the Dutch Ministry of Agriculture, Nature and Food Quality (Knowledge Base projects in the KB21 and KB34 program) and Top Sector Agri & Food (TKI-AF-15261).

